# Optimized methods for mapping DNA double-strand-break ends and resection tracts and application to meiotic recombination in mouse spermatocytes

**DOI:** 10.1101/2024.08.10.606181

**Authors:** Soonjoung Kim, Shintaro Yamada, Kaku Maekawa, Scott Keeney

## Abstract

DNA double-strand breaks (DSBs) made by SPO11 protein initiate homologous recombination during meiosis. Subsequent to DNA strand breakage, endo- and exo-nucleases process the DNA ends to resect the strands whose 5’ termini are at the DSB, generating long 3’-terminal single-stranded tails that serve as substrates for strand exchange proteins. DSB resection is essential for meiotic recombination, but a detailed understanding of its molecular mechanism is currently lacking. Genomic approaches to mapping DSBs and resection endpoints, e.g., S1-sequencing (S1-seq) and similar methods, play a critical role in studies of meiotic DSB processing. In these methods, nuclease S1 or other enzymes that specifically degrade ssDNA are used to trim resected DSBs, allowing capture and sequencing of the ends of resection tracts. Here, we present optimization of S1-seq that improves its signal:noise ratio and allows its application to analysis of spermatocyte meiosis in adult mice. Furthermore, quantitative features of meiotic resection are evaluated for reproducibility, and we suggest approaches for analysis and interpretation of S1-seq data. We also compare S1-seq to variants that use exonuclease T and/ or exonuclease VII from *Escherichia coli* instead of nuclease S1. Detailed step-by-step protocols and suggestions for troubleshooting are provided.

## Introduction

Meiosis is a specialized cell division used by sexually reproducing organisms to reduce the genome complement by half during gametogenesis. In this process, progenitor cells divide their genomic contents two times after one round of DNA replication (Hunter 2015). During the prophase of the first meiotic division, programmed DNA double-strand breaks (DSBs) are formed and repaired to produce physical links between homologous chromosomes, which ensure the faithful segregation of homologs in many species.

DSB formation is executed by a dimer of the evolutionarily conserved SPO11 protein, which uses a topoisomerase-like transesterase reaction to cleave the DNA strands and generate a pair of covalent protein-DNA complexes (Bergerat et al. 1997; Keeney et al. 1997) (**Figure 1A**). Each SPO11-bound DNA end is then digested by a combination of nucleases in a process known as DNA end resection, releasing a short DNA fragment still covalently bound to SPO11 (known as a SPO11-oligo complex) and exposing a long 3’-single-stranded DNA (ssDNA) tail that can subsequently initiate strand invasion for homologous recombination (**Figure 1A**) (Symington 2016; Cejka and Symington 2021). The outlines of the resection reaction are conserved between budding yeast and mice, but the molecular details of DSB processing remain incompletely understood, particularly in mammalian meiosis. Addressing this issue requires tools for examining the complexity of DSB processing genome wide under physiological conditions.

**Figure 1.**
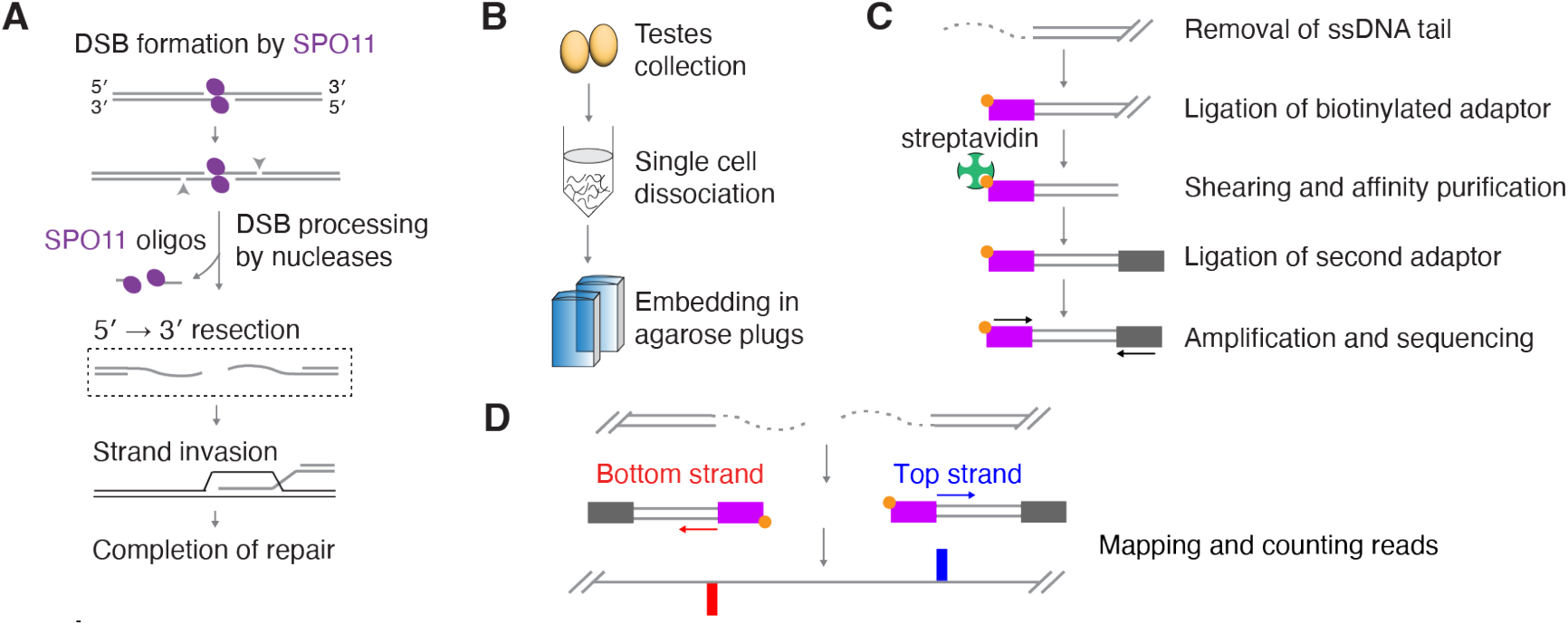
S1-seq overview. (A) DSB formation and processing in meiotic recombination. SPO11 (magenta ellipses) makes a DSB and remains covalently attached to 5’ strand termini. The SPO11-bound strands are subsequently nicked (arrowheads), and bi-directional resection by various nucleases releases SPO11-oligo complexes and exposes long 3’ ssDNA tails, which are used for homology search and strand invasion. (B) Preparation of DNA samples from whole testes. (C) Schematic of steps in S1-seq library preparation. (D) Schematic of mapping and counting the endpoints of S1-seq reads.

Recent years have seen the development of numerous whole-genome sequencing techniques aimed at visualizing the locations of DSBs and repair intermediates (Amente et al. 2021). To understand meiotic DSB resection, we developed the “S1-seq” method to visualize the global distribution of endpoints of meiotic DSB processing in yeast (Mimitou et al. 2017) and subsequently applied the method to mice (Yamada et al. 2020). Here, we provide an optimized S1-seq protocol for mouse testis samples that improves signal:noise ratio and allows application to samples from adult mice. We also explore which features of S1-seq maps are highly reproducible and which are less so. This reproducibility informs approaches to analysis and interpretation of S1-seq data. We further provide side-by-side comparison of S1-seq and related methods that use other nuclease cocktails to remove ssDNA tails for sequencing library generation, highlighting complementary strengths of the different methods.

## Results and Discussion

### Overview of S1-seq

To make S1-seq libraries from mouse testis samples, spermatocytes are first dissociated into single cells that are then embedded in low-melting-point agarose plugs to minimize random physical breakage of the genomic DNA. This ensures that only pre-existing DSBs that were generated in vivo are detected, while other genomic DNA remains largely intact during downstream steps (**Figure 1B**). While still in the plugs, cells are lysed and then DNA is deproteinated through proteinase K treatment, and the single-stranded DNA (ssDNA), a product of DSB end resection, is digested with nuclease S1. The blunted DSB ends are then polished and ligated to biotinylated sequencing adaptors (**Figure 1C**). Genomic DNA is subsequently extracted from the agarose, randomly sheared by sonication, and the fragments of DNA that were ligated to the first adaptor are enriched by affinity purification with streptavidin. A second, DSB-distal adaptor is then ligated to the sheared ends and the DNA fragments are amplified by PCR with indexed primers and sequenced (**Figure 1C**). The sequencing reads are mapped and the resection endpoint is assigned at the nucleotide next to the biotinylated adaptor sequence (**Figure 1D**).

Upon mapping to an appropriate mouse genome assembly, S1-seq reads cluster around meiotic DSB hotspots (**Figure 2A**) (Yamada et al. 2020). Moreover, the reads are distributed around hotspot centers with defined polarity: top-strand reads are enriched to the right because they arise from resection tracts moving away from DSBs to the right, while bottom-strand reads reflect tracts moving to the left (Mimitou et al. 2017;

**Figure 2.**
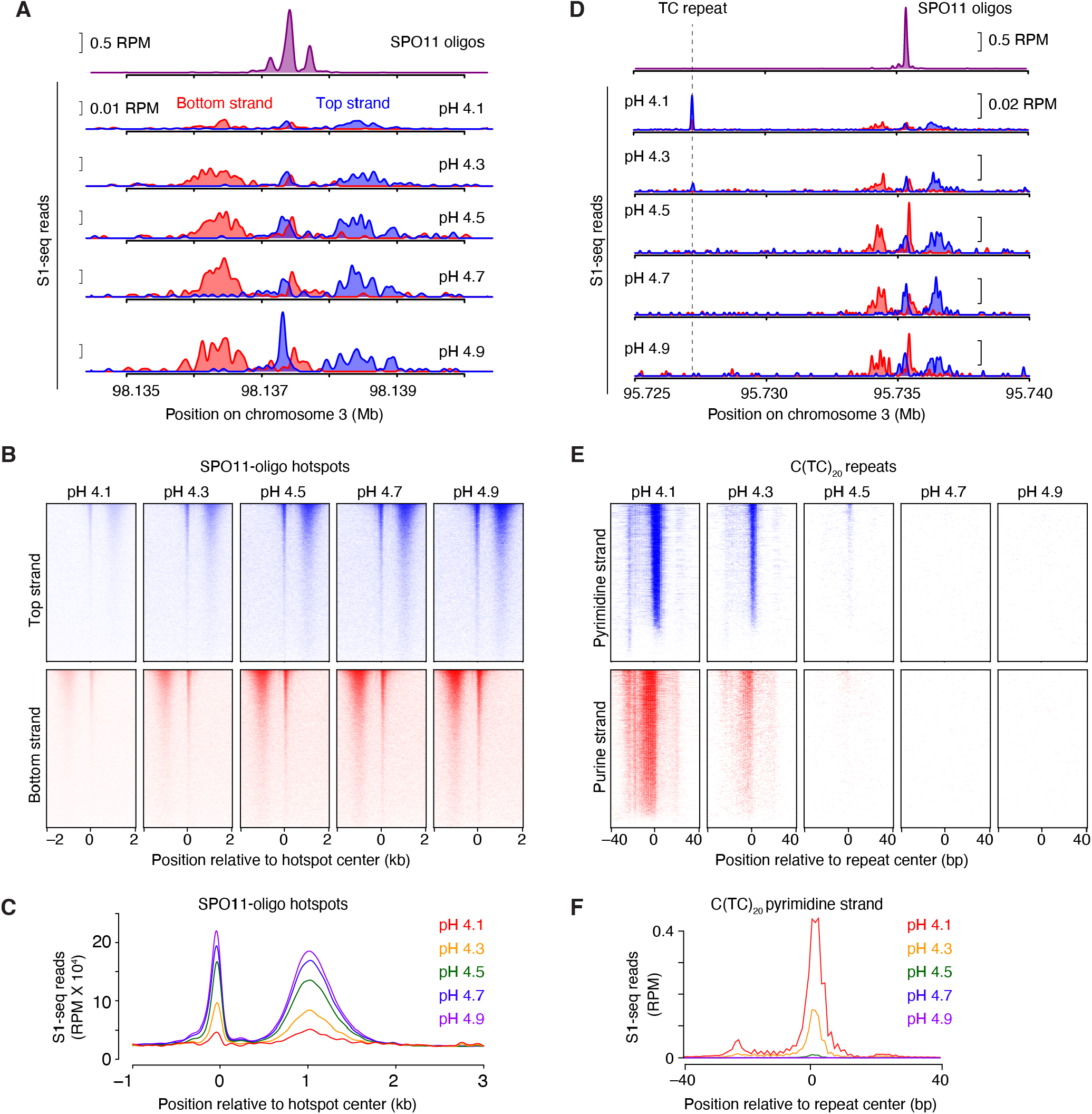
Effect of pH on S1-seq detection of non-B-form DNA. S1-seq was performed on genomic DNA isolated from whole testes of 14.5-dpp mice. For each map, the nuclease S1 digestion step was carried out at the indicated pH. Panels A–C show S1-seq reads around meiotic DSB hotspots; panels D–F show reads around representative H-DNA-forming TC repeats. (A) Strand-specific S1-seq signals at a representative DSB hotspot, smoothed with a 151-bp Hann window. SPO11-oligo sequencing data shown here (top plot) and throughout are from Lange et al. (2016). The baseline of the y axis for each plot is 0. (B) Heatmaps (data in 50-bp bins) of strand-specific S1-seq reads around SPO11-dependent DSB hotspots from wild type [n = 13,960 hotspots from Lange et al. (2016)]. Each line is a hotspot, ranked from strongest at the top. (C) Genome-average profiles of S1-seq reads around SPO11-oligo hotspots. The bottom-strand reads were reoriented and combined with the top-strand reads, averaged across all hotspots, then the signal was smoothed with a 151-bp Hann window. (D) pH dependence of S1-seq signal at a representative H-DNA-forming TC repeat. Note that the signal at the TC repeat becomes weaker with increasing pH, while the signal becomes stronger (i.e., better signal:noise ratio) for the SPO11-dependent hotspot nearby. (E) pH dependence of S1-seq signal intensities around C(TC)20 repeats [n = 2,664 (Maekawa et al. 2022)]. Heatmaps are plotted without binning. Reads mapping to the pyrimidine strand are shown in blue; purine-strand reads are in red. (F) Genome-average profiles of S1-seq reads on the pyrimidine strand at C(TC)20 repeats.

Mimitou and Keeney 2018; Yamada et al. 2020) (**Figures 1D and 2A**). There is also a prominent signal near hotspot centers that is thought to arise from S1 cleavage of recombination intermediates (Yamada et al. 2020), discussed further below.

### Optimized pH of the S1 digestion step improves signal:noise ratio for meiotic DSBs

We recently reported that a major source of SPO11-independent S1-seq signal in mouse genomic DNA reflects nuclease S1 cleavage at H-DNA motifs (Maekawa et al. 2022), particularly polypyrimidine•polypurine mirror repeats such as C(TC)_20_, which can form an intramolecular triplex plus ssDNA (Mirkin and Frank-Kamenetskii 1994)p. We considered it likely that the H-DNA triplex formation we detected might be occurring *ex vivo* during sample handling because nuclease S1 activity is optimal under acidic conditions that also favor the formation of precisely the type of H-DNA we observed (Mirkin and Frank-Kamenetskii 1994)p.

To test this idea, we varied the pH of the S1 digestion step. For this experiment, the sequencing libraries were generated from plugs made from the same testis cell preparation in the same reaction batch, so the amount of SPO11-generated DSBs should be the same in every sample. However, the final S1-seq signals at hotspots, normalized per million mapped reads (RPM), showed marked differences between the maps. At the lowest values tested (pH 4.1 or 4.3), S1-seq signals around SPO11-dependent DSB hotpots were detectable but weak (**Figure 2A– C**). The weakness of the signal reflects the fact that a large fraction of total reads maps to H-DNA motifs instead (**Figure 2D–F**) (Maekawa et al. 2022). This depresses the calculated amount of SPO11-dependent signal when normalized to the total number of mapped reads and, more importantly, also decreases the sequencing read depth at hotspots, which increases the noise from sampling error.

The S1-seq signal at H-DNA motifs was exquisitely pH-sensitive, becoming extremely weak at pH 4.5 and undetectable at pH 4.7 and 4.9 (**Figure 2D–F**). These findings confirm our hypothesis that most or all of the H-DNA detected by S1-seq in our experiments arose *ex vivo* during sample preparation (Maekawa et al. 2022)/. By removing this artifactual background, meiotic DSB hotspots showed clear improvement of signal:noise ratio (**Figure 2A–C**).

The S1-seq patterns around hotspots are highly stereotyped, showing very similar spatial distributions at every hotspot (Yamada et al. 2020) (**Figure 2B**). Therefore, we routinely derive a global S1-seq signature by co-orienting and combining the sequencing signal from top and bottom strand reads and averaging across all hotspots (**Figure 2C**). These average profiles facilitate comparisons between different datasets. To illustrate, we overlaid the average profiles from S1-seq at pH 4.3 and pH 4.7 with and without smoothing (**Figure 3A**). Smoothing improves the signal visualization without distorting the overall spatial distribution.

**Figure 3.**
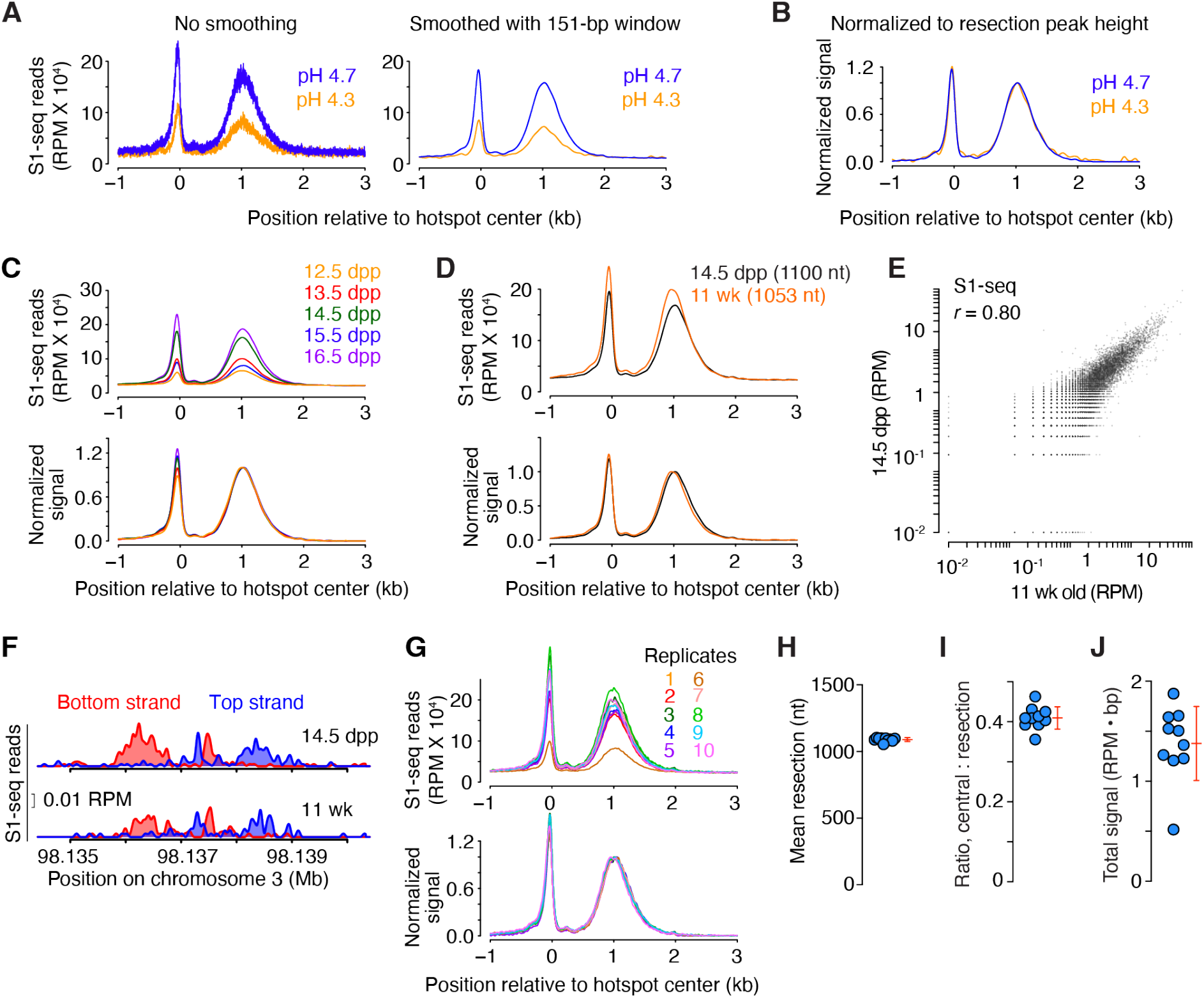
Reproducibility of S1-seq maps. (A,B) Smoothing and internal normalization for display of genome-wide average resection profiles. In panel A, the average S1-seq reads at SPO11-dependent hotspots generated by S1 digestion at either pH 4.3 or pH 4.7 (from Figure 2C) are presented again without (left) or with smoothing with a 151-bp Hann window. In panel B, the smoothed average profiles were internally normalized to set the height of the resection peak to a value of one after subtracting background (defined as the value of the signal 2,500 bp to the right of the hotspot center). Negative values after background subtraction were set as zero for plotting purposes. (C) Reproducibility of S1-seq spatial patterns but not signal strength at various juvenile time points. Genome-average profiles around hotspots are shown for S1-seq maps generated from whole testes of mice of the indicated ages. Samples were collected from a single litter daily at approximately the same time each day. Plugs were prepared immediately after harvest and stored until the final time point, then S1-seq libraries were generated in a single batch. Internally normalized (bottom) and non-normalized (top) profiles are shown. (D) Similar genome-average S1-seq profiles from juvenile (14.5 dpp) and adult (11 wk old) testis samples. Internally normalized (bottom) and non-normalized (top) profiles are shown. Values in parentheses are the mean resection lengths. To calculate these means, S1-seq sequencing signal was averaged across hotspots and an estimated background was removed by subtracting from all values the value of signal 2.5 kb away from the hot spot center. The signal close to and further away from the hot spot center was excluded by setting values of positions <100 bp and >2.5 kb to zero. Fractions of total signal were calculated every 100 bp and the mean resection length was calculated. (E) Correlation (Pearson’s r) of S1-seq read counts from juvenile and adult samples. Each point is a SPO11-oligo hotspot, with S1-seq signal summed from −2000 to −250 bp (bottom strand) and +250 to +2000 bp (top strand) relative to the hotspot center. Hotspots with no signal (0 RPM) were excluded for Pearson’s r calculation. Hotspots with <10−2 RPM were set as 10−2 for plotting purposes. (F) Strand-specific S1-seq at a representative DSB hotspot. (G) Reproducibility of S1-seq profiles around hotspots in biological and technical replicates. Internally normalized (bottom) and non-normalized (top) profiles are shown. A smaller smoothing window (51-bp Hann window) was used here to highlight small differences between libraries. Replicate #1 is the 14.5-dpp sample from panel D; #2 is the 14.5-dpp sample from panel C; #4 is a technical replicate of #1; #5 is the pH 4.7 sample from panel A; #10 is the adult sample from panel D; and the other profiles are additional biological replicates from 14.5-dpp (#3), 15.5-dpp (# 6, 8), and 16.5-dpp (#7, 9) samples not shown elsewhere. (H–J) Reproducibility of quantitative features of S1-seq genome-average profiles around hotspots. Each point represents one of the replicates shown in panel G for mean resection length (H); the ratio of the central peak (summed from –1500 to +100 bp) to the resection peak (summed from +100 to +2500 bp) (I); and total signal summed from –1500 to +2500 (J). Error bars indicate mean ± SD.

Because it is difficult to evaluate by eye the similarity between the shapes of two curves of different signal strength (**Figure 3A**), we further internally normalize the average profiles by setting the height of the resection peak to a value of 1 (**Figure 3B**). Importantly, even though the signal levels differed starkly between the pH 4.3 and pH 4.7 samples, their normalized spatial distributions were virtually identical (**Figure 3B**). Therefore, when comparing multiple S1-seq libraries, we suggest using the internally normalized pattern for a rigorous evaluation of potential differences in resection endpoint distributions.

### Spermatocytes from juvenile and adult mice show highly similar resection patterns

Initially, we developed S1-seq for *Saccharomyces cerevisiae* meiotic cells under synchronous culture conditions (Mimitou et al. 2017; Mimitou and Keeney 2018). Our first attempts to generate S1-seq maps from whole testes of wild-type adult mice were unsuccessful, presumably because the low signal:noise ratio with S1-seq performed at pH < 4.5 was exacerbated by the lack of meiotic synchrony and because spermatocytes are only a subset of the cells present in adult testis tissue (S. Yamada and S. Keeney, unpublished observations).

To overcome these issues, we previously took advantage of the semi-synchronous first wave of spermatogenesis in juvenile mice (Yamada et al. 2020). Meiotic DSB formation and repair first occur in the juvenile testis between 9 and 16 days postpartum (dpp), at which time later stage germ cells (e.g., round and elongating spermatids) are not present (Bellve et al. 1977; Guillon et al. 2005; Lange et al. 2011; Zelazowski et al. 2017). Using testes of juvenile mice thus enriches for early meiotic prophase I cells, which enabled us to obtain S1-seq datasets with good signal:noise ratio (Yamada et al. 2020).

To assess whether meiotic resection patterns change during prophase I, we generated S1-seq maps from mice ranging in age from 12.5 to 16.5 dpp (**Figure 3C**). The spatial distribution of resection endpoints was remarkably uniform across this time span (**Figure 3C bottom**), despite variations in signal intensity prior to internal normalization (**Figure 3C top**). Because the testes of mice at these ages differ substantially with respect to which substages of prophase I are prevalent (Bellve et al. 1977), we infer that resection is completed rapidly after DSB formation and that each resection tract is then stably maintained, similar to the situation in *S. cerevisiae* (Mimitou et al. 2017).

One downside to using juvenile mice is that it requires greater care in monitoring birth timing and allows only a short window of a few days for sample collection. Inability to assay adult tissues also hinders the broader application of S1-seq to other multicellular species that do not have the same potential to harvest naturally synchronized germ cell populations. The optimized protocol presented here overcame these limitations and allowed us to successfully perform S1-seq on samples from adult mice. Datasets from adults were highly similar to those from juveniles for average DSB resection lengths (**Figure 3D**), hotspot strength (as measured by S1-seq read counts; **Figure 3E**), and local patterns around individual hotspots (**Figure 3F**). These similarities show that findings from juvenile mice can be directly extrapolated to meiosis in adults, which is important because some aspects of meiotic recombination operate differently during the juvenile spermatogenesis waves than in adults (Zelazowski et al. 2017).

### Reproducibility of the quantitative features of S1-seq DSB maps

We sought to determine which aspects of S1-seq maps are highly reproducible and which are less so. To do this, we compared ten maps from both biological and technical replicate S1-seq experiments from wild-type juvenile or adult testes (**Figure 3G**). In this analysis, the spatial distribution of resection endpoints was extremely reproducible, with biological replicates giving highly superimposable curves across the region from ∼0.2 to 3 kb from hotspot centers (**Figure 3G**). As a result, estimates of average resection lengths varied little between samples (1089 ± 15 nt, mean ± SD, for a coefficient of variation (CV) of just 1.4%) (**Figure 3H**). The central signal was also highly reproducible, both in shape (**Figure 3G**) and signal strength (measured as area under the curve) relative to the resection signal (**Figure 3I**; CV = 6.8%).

By contrast, the absolute sequencing signal strength in RPM was less reproducible from sample to sample (**Figure 3J**; CV = 26.9%), indicating that the magnitude of the S1-seq signal from meiotic DSBs varies relative to other sources of sequencing reads. Possible causes of this variability in signal-to-noise ratio could include batch-to-batch variation in detection of non-B-form DNA structures (e.g., H-DNA, as described above) and sample-to-sample differences in cellular composition of the testis cell suspensions (e.g., ratio of spermatocytes to other cell types that contain ssDNA, e.g., from DNA replication). Regardless of the source, this variability means that less weight should be placed on changes in absolute signal strength unless those changes are well outside the range of variation observed in biological replicates.

### Alternatives to S1 nuclease for removal of ssDNA from resected DSB ends

Another method for mapping DSB ends, called END-seq, was developed by others independently (Canela et al. 2016). A principal difference between the methods is the nuclease used to remove ssDNA, with END-seq employing a combination of *E. coli* exonuclease VII and exonuclease T (Canela et al. 2016; Canela et al. 2017; Paiano et al. 2020; Wong et al. 2021). We adapted our S1-seq protocol to similarly use a combination of exonuclease VII and exonuclease T (referred to hereafter as Exo7/T-seq) or exonuclease T alone (termed ExoT-seq).

The distinct substrate specificities of each enzyme yield both overlapping and contrasting results. All three enzymes perform 3’ ssDNA digestion, resulting in blunted DNA ends suitable for adaptor ligation (although often needing polishing with T4 DNA polymerase) (**Figure 4**). However, among the three, only nuclease S1 possesses endonuclease activity capable of digesting recombination intermediates such as D loops (**Figure 4A**). This unique feature gives a distinctive central signal in S1-seq: reads are offset slightly from the hotspot centers, with opposite strand polarity to that seen for resected DSB ends, i.e., top strand reads clustering to the left of hotspot centers rather than being on the right side (Mimitou et al. 2017; Mimitou and Keeney 2018; Yamada et al. 2020) (**Figure 4A**). This central signal is thought to arise from interhomolog recombination intermediates because it is absent in animals lacking the strand-exchange protein DMC1 and is not seen at hotspots on the non-homologous parts of the X and Y chromosomes in wild type (Yamada et al. 2020).

**Figure 4.**
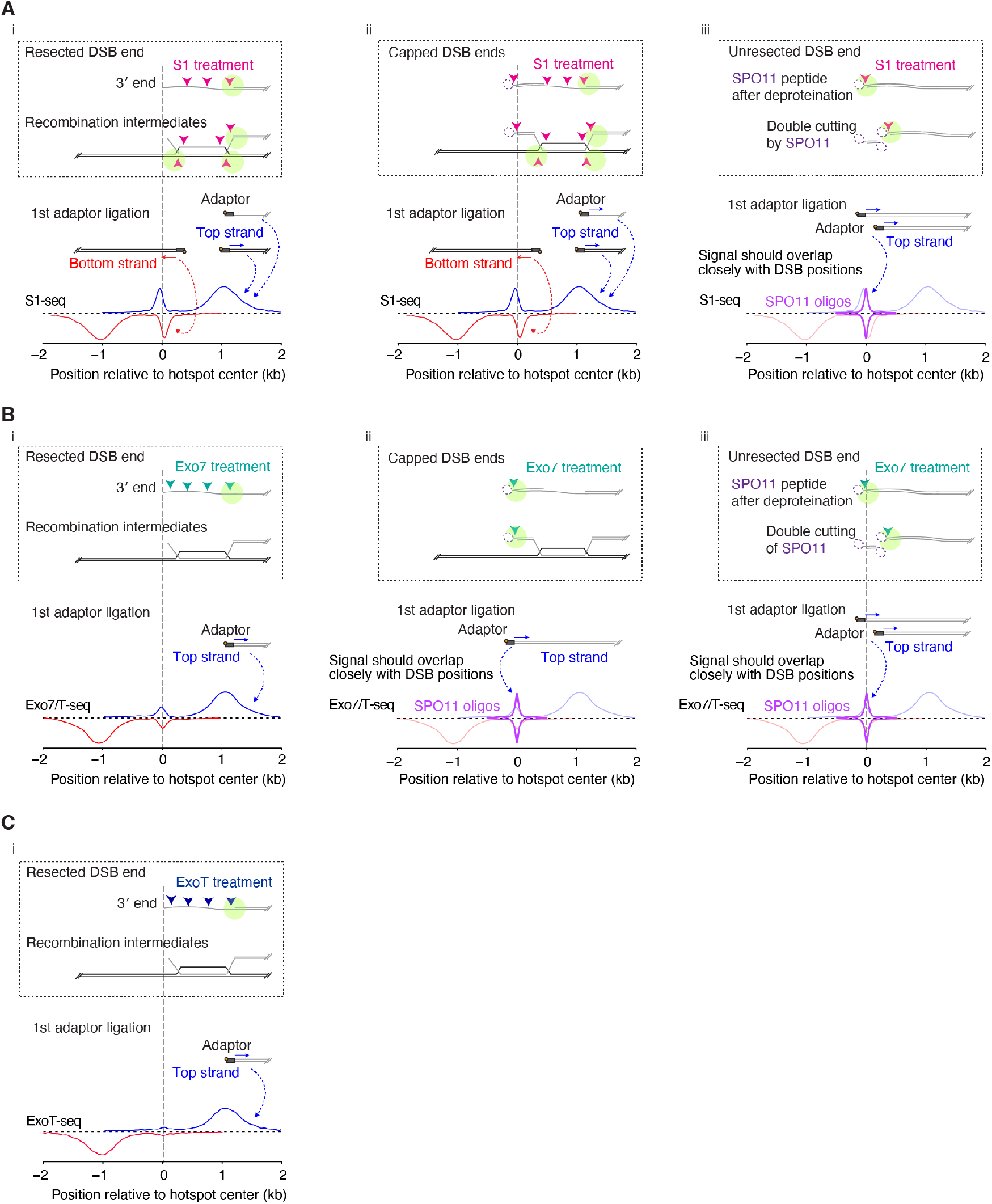
Presumptive processing of DSBs and other meiotic recombination intermediates by the nuclease activities of S1-seq, Exo7/T-seq, and ExoT-seq. The schematics illustrate how each nuclease cleaves free, resected DSB ends (i); SPO11-capped resected ends (ii); or unresected DSBs (iii) plus relevant strand-exchange intermediates (i and ii). The vertical dashed lines align the different elements of the cartoons by the 3’ end of the DSB ssDNA tail. SPO11 cuts DNA with a 2-nt 5’ overhang and stays covalently bound to the over-hang until released from DSB ends by resection (Liu et al. 1995; Pan et al. 2011). The existence of a SPO11-bound (capped) recombination intermediate was proposed previously (Paiano et al. 2020; Yamada et al. 2020). During library preparation, deproteination by proteinase K removes the DNA-bound SPO11 protein from capped or unresected ends, leaving a small peptide adduct (dashed magenta ellipses). Each panel depicts how digestion of the indicated structures (arrowheads) by nuclease S1 (A), exonuclease VII (B), or exonuclease T (C) could generate duplex DNA ends suitable for sequencing adaptor ligation (pale green circles). (A) Cleavage by S1 endonuclease activity of ssDNA from resected, capped, and unresected breaks in addition to D-loop recombination intermediates. During later steps of library preparation, the short duplex oligonucleotides produced from capped ends and D-loops are not recovered (Yamada et al. 2020). Therefore, S1-seq primarily generates reads from resection endpoints, which are distributed approximately 1 kb away from the center of hotspots with the expected polarity, and from recombination intermediates near the hotspot center (referred to here as the central signal) that have opposite polarity. (B) Presumed cleavage by exonuclease VII of resected, capped, or unresected breaks and recombination intermediates. Exonuclease VII is a ssDNA-specific exonuclease that degrades overhangs in either the 3’-to-5’ or 5’-to-3’ direction from a DNA terminus, producing 4–10 nt oligonucleotides as a result (Chase and Richardson 1974a; Chase and Richardson 1974b; Liu et al. 2024). Exonuclease VII can also remove peptide blocks such as SPO11 or aborted topoisomerase II from DNA ends after proteinase K digestion and can generate blunt DNA ends when combined with exonuclease T in the END-seq procedure (Paiano et al. 2020; Huang et al. 2021; Wong et al. 2021). Neither exonuclease VII nor exonuclease T is expected to be able to digest D-loop structures, so it is likely that the central signal from Exo7/T-seq, which overlaps closely with hotspot centers, is derived from capped or unresected DSB ends (Paiano et al. 2020). (C) Presumed cleavage by exonuclease T of resected DSB ends or recombination intermediates. Exonuclease T catalyzes the removal of nucleotides from linear ssDNA or RNA in the 3’-to-5’ direction (Zuo and Deutscher 1999). Exonuclease T is not able to degrade blocked DNA ends or D-loops, so ExoT-seq provides only resection endpoint signals and little or no central signals.

Exonuclease VII can remove 5’ overhangs, including those blocked by protein adducts (Chase and Richardson 1974a; Huang et al. 2021; Liu et al. 2024). Consequently, the central signal seen in END-seq (or Exo7/T-seq) is thought to arise from a subset of DSB ends that remain capped by SPO11-oligo complexes bound to the two-nucleotide 5’ overhang left by the SPO11 DNA-cleavage activity (**Figure 4B**) (Paiano et al. 2020; Yamada et al. 2020). Additionally, both S1-seq and Exo7/T-seq are able to generate sequencing signals from DSBs that have not undergone any resection at all (**Figures 4A**,**B**) (Mimitou et al. 2017; Mimitou and Keeney 2018; Paiano et al. 2020; Yamada et al. 2020; Claeys Bouuaert et al. 2021). Detection of unresected DSBs in END-seq appears to be inefficient, but can be improved by treatment of the DNA with tyrosyl DNA phosphodiesterase 2 (TDP2) (Paiano et al. 2020), which can remove the fragments of SPO11 that remain covalently bound to DNA after proteolysis (Gittens et al. 2019). The central signals from capped or unresected DSB ends are closely congruent with DSB positions defined by SPO11-oligo sequencing (**Figures 4A**,**B**).

In contrast to the other methods, ExoT-seq can only detect cleanly resected DSBs because exonuclease T requires a single-stranded DNA end with a free 3’-OH (Deutscher and Marlor 1985). Consequently, Ex-oT-seq maps almost completely lack the central signal (**Figure 4C**).

Figure 5. compares results from S1-seq, Exo7/T-seq, and ExoT-seq. All three methods gave essentially indistinguishable spatial patterns for resected DSB ends, with highly congruent measurements of mean resection lengths (**Figures 5A–C**). The major difference is the signal at the hotspot center, which was shifted to the left of hotspot midpoints in S1-seq, weaker and centered on hotspot midpoints in Exo7/T-seq, and essentially absent in ExoT-seq (**Figure 5C**). As with S1-seq, results were virtually identical when comparing Exo7/T-seq between juveniles and adults (**Figure 5D**).

Using Exo7/T-seq, we also tested whether sequencing libraries could be generated from stored testis samples. We compared results generated from cells embedded in agarose immediately after animal sacrifice and testis harvest with results obtained by embedding cells after storage of testes on ice for 1, 3 or 5 days after harvest, or after slow freezing of testes in fetal bovine serum containing 10% DMSO. The sample stored for 1 day on ice was nearly indistinguishable from the freshly prepared sample, but meiotic DSB hotspot-proximal signals from the samples stored longer on ice or after freezing were significantly reduced (**Figure 5E, top**). The reduced signal:noise ratio from the latter samples may be due in part to an increased fraction of inviable cells and a decreased efficiency of single cell dissociation (**Figure 5F**), contributing to background signals from randomly broken DNA ends through DNA fragmentation. Notably, however, even under conditions with weaker signal:noise ratios, the global averages for both the resection and central signals were indistinguishable from fresh samples after internal normalization (**Figure 5E, bottom**). We conclude that the agarose plugs should ideally be prepared with tissue that is freshly harvested or stored no longer than about one day on ice, but that it is still feasible to analyze DSB resection patterns from testis samples that are stored longer or even frozen if necessary.

**Figure 5.**
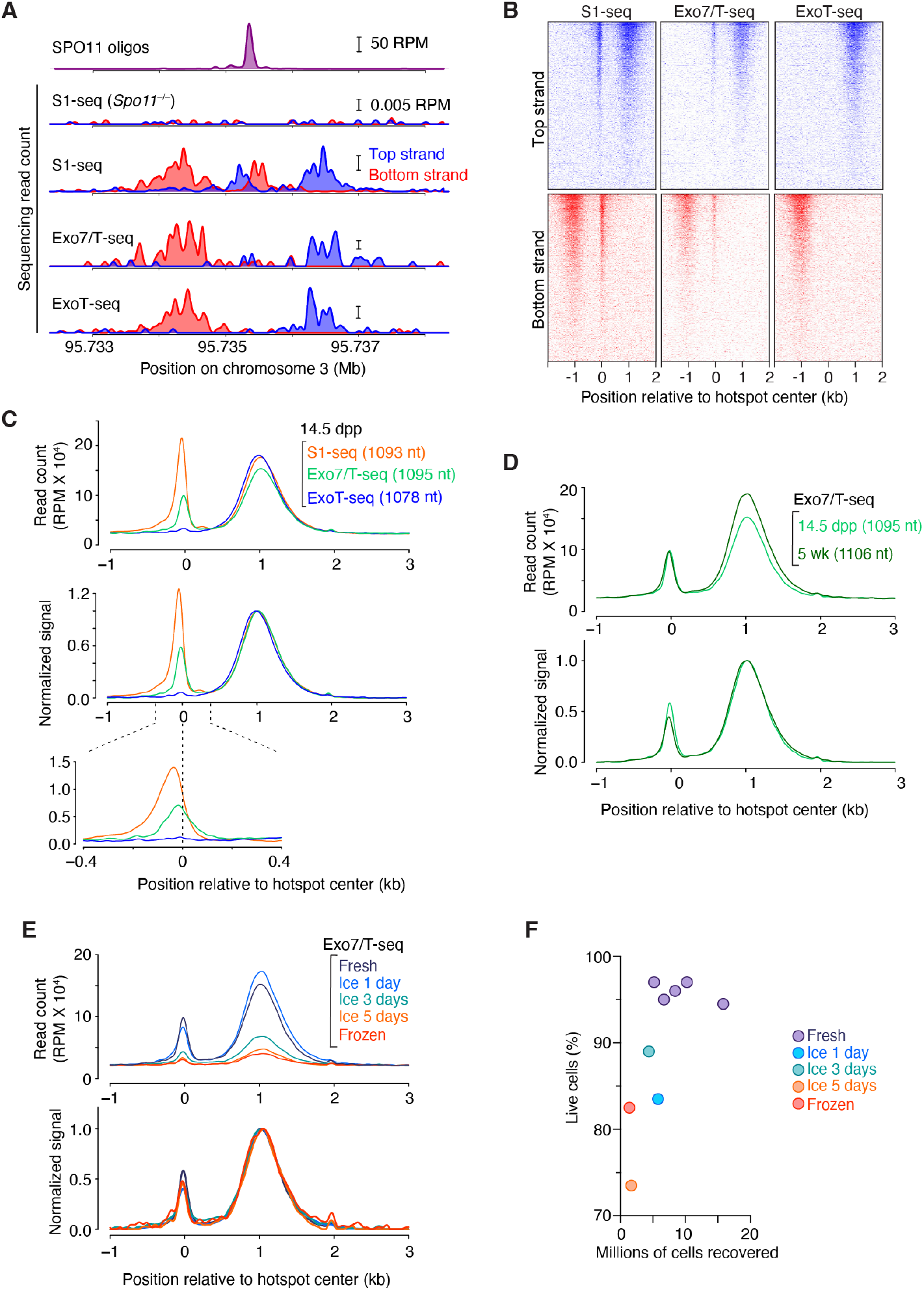
Comparisons of results from S1-seq, Exo7/T and ExoT-seq methods. (A) Strand-specific resection endpoint signals mapped by S1-seq, Exo7/T-seq or ExoT-seq at a representative DSB hotspot in 14.5-dpp mice. Signals from three (S1-seq for wild type) or two (other methods and S1-seq for Spo11–/–) biological replicate libraries were averaged. Data are smoothed with a 151-bp Hann window. (B) Stereotyped distribution of resection endpoints and central signal around DSB hotspots. One representative library is shown for S1-seq and two each are averaged for the others. Each line is a SPO11-oligo hotspot (n = 13,960), strongest at the top. Sequencing signals (in 40-bp bins) were locally normalized by dividing by the total signal in a 4001-bp window around each hotspot’s center. Each hotspot thus has a total value of 1, so that spatial patterns can be compared between hotspots of different strengths. (C) Genome-average profiles of S1-, Exo7/T- or ExoT-seq reads from 14.5-dpp mice. Signals from three (S1-seq) or two (other methods) biological replicate libraries were averaged. Values in parentheses indicate mean resection lengths calculated as described in Figure 3D. Non-normalized (top) and internally normalized (middle) profiles are smoothed with a 151-bp Hann window. The bottom-most panel shows zoomed view (smoothed with a 51-bp Hann window) into the region around hotspot centers. (D) Comparison of genome-average Exo7/T-seq profiles from 14.5-dpp juvenile or 5-wk-old adult mice. The 14.5-dpp profile is reproduced from panel C. Two biological replicate libraries were averaged for each age. Mean resection lengths are indicated in parentheses. (E) Genome-average Exo7/T-seq profiles generated from cells embedded in agarose from fresh testes immediately after harvest, or from testes that were either stored on ice or frozen. The profile for the fresh preparation is reproduced from panel C. (F) The number of cells recovered and percentages of viable cells after single-cell dissociation from fresh testes or testes stored as indicated. In C, D, and E, internally normalized (bottom) and non-normalized (top) profiles are shown.

### Detailed protocols

#### 1 Mouse testis cell dissociation and plug preparation

##### 1.1 Key equipment and consumables

- Low-retention filter tips
- 1.5 ml low surface adhesion ubes
- 15 ml and 50 ml conical tubes
- 14 ml round bottom tubes (Falcon, cat. no. 352059)
- 70-μm cell strainer (Falcon, cat. no. 352350)
- Transfer pipet (Sigma, cat. no. Z350605)
- 50-well disposable plug molds (Bio-Rad, cat. no. 1703713)
- Paraffin film
- Dumont forceps super fine #5
- Tabletop centrifuge
- Thermal mixer (e.g., Eppendorf ThermoMixer C)
- Heating block
- Hybridization oven
- 0.22 μm cellulose nitrate sterilizing filter
- Automatic cell counter or hemocytometer

##### 1.2 Reagents and buffers

- PBS without Ca^2+^ and Mg^2+^
- Dulbecco’s Modified Eagle Medium (DMEM) (optional: without phenol red for visibility)
- Fetal bovine (or calf) serum
- UltraPure DNase/RNase-free distilled water (Thermo Fisher Scientific, cat. no. 10977015)
- Gey’s balanced salt solution (GBSS) (Sigma, cat. no. G9779)
- Dispase II (Sigma, cat. no. D4693); prepare 10 mg/ml stock solution with PBS without Ca^2+^ and Mg^2+^ and store aliquots at –80 °C.
- Collagenase IV (Worthington Biochemical, cat. no. LS004186); prepare approximately 10 mg/ml stock solution by add 9 ml of GBSS to 100 mg powder and store aliquots at –80 °C.
- DNase I (Roche, cat. no. 10104159001); prepare 1 mg/ ml stock solution with UltraPure DNase/RNase-free distilled water and store aliquots at –80 °C.
- Polyvinyl alcohol (PVA) (Sigma, cat. no. P8136); prepare 10% w/v solution with UltraPure DNase/RNase-free distilled water and dissolve by autoclaving and store at 4 °C.
- Bovine albumin fraction V (7.5% solution) (Thermo Fisher Scientific, cat. no. 15260037)
- TrypLE Select Enzyme (1X), no phenol red (Thermo Fisher Scientific, cat. no. 12563011)
- Low-melting point SeaPlaque GTG Agarose (Lonza, cat. no. 50115); using a boiling water bath, prepare 1% w/v solution with GBSS and store 1 ml aliquots at 4 °C until use. Low-melting point agarose from Bio-Rad (cat. no. 1613111) is an acceptable alternative, but we have had difficulty generating libraries using low-melt agarose from Invitrogen (cat. no. 16520050).
- Proteinase K (Roche, cat. no. 3115879001); reconstitute according to manufacturer’s instruction.
- RNase A (Thermo Fisher Scientific, cat. no. EN0531)
- 0.5 M EDTA pH 8.0
- 1 M Tris-HCl pH 7.5
- N-lauroylsarcosine sodium salt (Sigma, cat. no. L9150); prepare 10% w/v solution with UltraPure DNase/ RNase-free distilled water, filter sterilize and store at 4 °C.
- TE buffer: 10 mM Tris-HCl at pH 7.5, 1 mM EDTA at pH 8.0, filter sterilize and store at room temperature.
- Lysis buffer: 0.5 M EDTA at pH 8.0, 1% N-lauroylsarcosine sodium salt, prepare freshly before use.

##### 1.3 Procedures

###### 1.3.1 Cell dissociation from mouse testes

1. Euthanize the mouse and remove the testes. Tear and pull the tunica albuginea with forceps to decapsulate the testes. Transfer both testes to a 50 ml conical tube with 3 ml of DMEM containing 0.1% PVA and 0.1% BSA (pre-warmed to 35 °C). Add 1 mg/ml of dispase II and 1 mg/ml of collagenase type IV. To guarantee the same length of enzyme treatment when processing multiple samples, add the collagenase and dispase after each sample has been transferred to a 50 ml conical tube.
2. Incubate for 20 minutes at 35 °C in an Eppendorf thermomixer at 450 rpm. Invert the tube gently every 5 min. The testes should break into discrete, short tubule segments; if the tubules are under-or over-dissociated, adjust the shaking speed or duration of incubation to reach the ideal state of dissociation (**Figure 6A**). **<u><u>Note 1</u></u>**: If an Eppendorf thermomixer is not available, it is okay to use a shaking incubator at 100 rpm or a slower speed. If shaking is too vigorous, over-dissociation of seminiferous tubule is often observed, which causes a poor yield of spermatocytes.
3. Collect dispersed seminiferous tubules by allowing them to settle to the bottom of the tube (**≥** 2 min). Remove the supernatant carefully with a transfer pipet.
4. Add 5 ml of DMEM containing 0.1% PVA, 0.1% BSA, and 1 μg/ml DNase I (pre-warmed 35 °C) to the tube, wash tubules by gentle inversion of the tube, and allow the tubules to settle for 2 min. Remove supernatant carefully with a transfer pipet.
5. Repeat step 4 twice (in total 3 washes).
6. Resuspend tubules in 3 ml of TrypLE Select Enzyme containing 0.1% PVA and 1 μg/ml DNase I. Incubate for 15 min at 35 °C in a thermomixer at 450 rpm, then add 150 μl FBS (final 5%) to inactivate TrypLE Select Enzyme.
7. Pipet up and down gently using a transfer pipette for 3 min to dissociate clumps of cells into single cells.
8. Add 5 ml of cold GBSS containing 0.1% PVA, 0.1% BSA.
9. Assemble a 70-μm cell strainer on top of a 50 ml conical tube and sieve the suspension through the strainer to remove clumps.
10. Transfer cells to a new 15 ml conical tube and place on ice.
11. Add 5 ml of cold GBSS containing 0.1% PVA, 0.1% BSA to the 50 ml tube in step 9 and rinse the inside of the tube to collect residual cells. Place on ice.
12. Spin cells in the 15 ml tube at 1,000 rpm (220 *g*) at 4 °C for 3 min.
13. Remove supernatant carefully with a transfer pipet and resuspend cells with the suspension from step 11 using a transfer pipet.
14. Repeat the wash two more times with 5 ml of cold GBSS containing 0.1% PVA, 0.1% BSA.
15. Prior to the last spin, take a small sample of the cell suspension and then count the cell number during the spin, using an automatic cell counter or a hemocytometer. Decide on the desired number of plugs. **<u>Note 2</u>:** Typically, 6∼8 million cells are obtained per juvenile mouse and 60∼80 million cells per adult mouse. For best performance in the following steps, prepare 1.5∼2 million cells per plug from a juvenile sample and 2.5∼3 million cells per plug from an adult sample. One plug from juvenile (1.5∼2 million cells) and two plugs from adult (5∼6 million cells) are typically enough to generate a successful library.
16. Prepare one 1.5 ml low surface adhesion tube per sample by rinsing the inside of the tube with GBSS containing 0.1% PVA, 0.1% BSA. Briefly spin the tubes and remove any residual liquid.
17. Resuspend cells with 1 ml of room temperature GBSS (without PVA or BSA) and transfer them to the 1.5 ml tube prepared in step 16. For this step, use 1 ml low-retention filter tips.
18. Spin cells at 1,000 rpm (220 *g*) at 4 °C for 2 min using a swinging bucket rotor. Using a swinging bucket rotor is essential to creating a tight pellet and avoiding cell loss.

**Figure 6.**
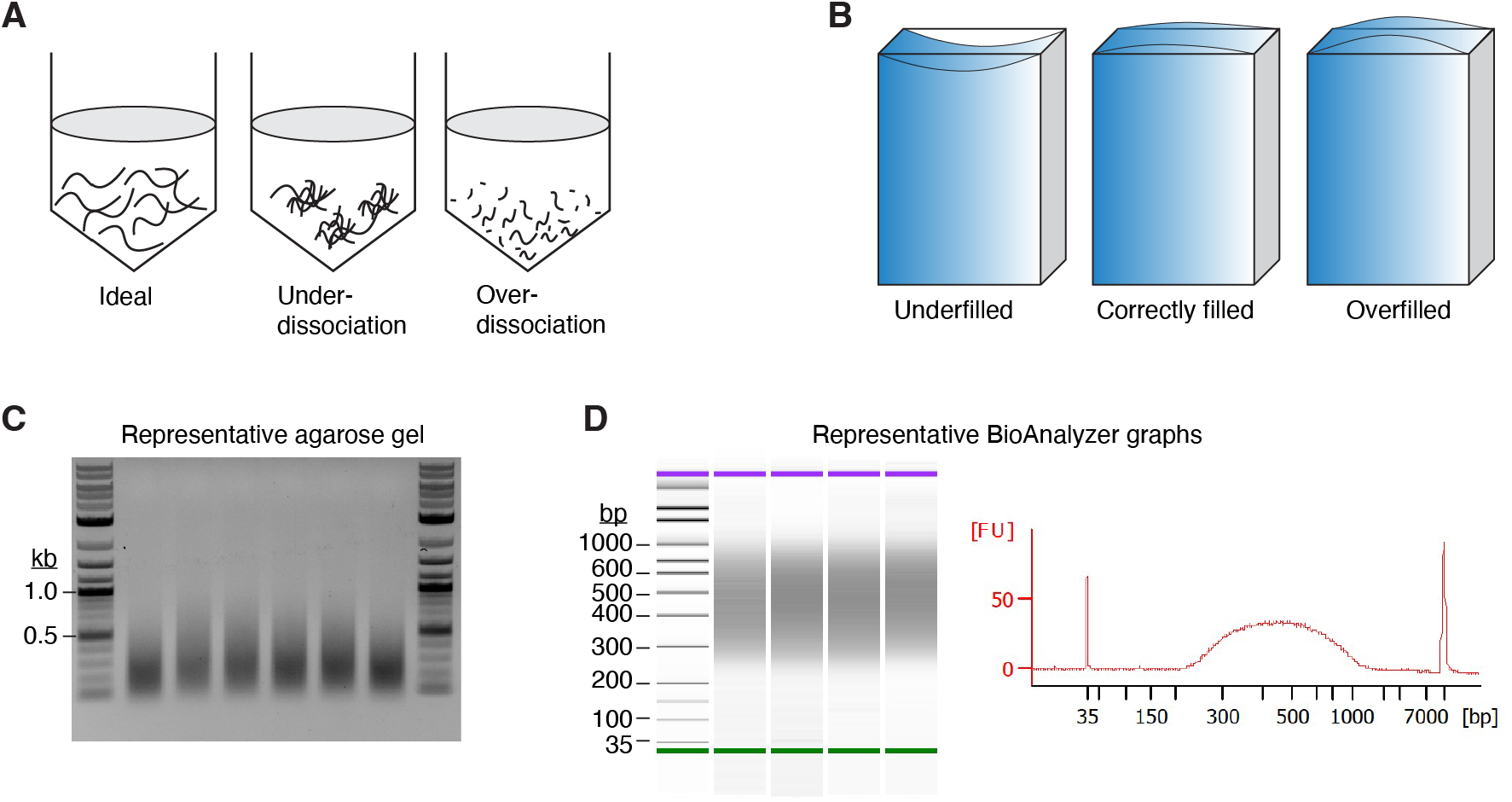
Procedural details and quality-control checks during sequencing library preparation. (A) Schematic showing the optimal dissociation of seminiferous tubules (left). If seminiferous tubules are under-dissociated (middle) or over-dissociated (right), the final cell count recovered from each testis will be smaller than expected. (B) Schematic showing the optimal level of agarose in a plug mold. (C) A representative gel image showing genomic DNA after sonication. About 1% of total DNA from each plug was loaded in each lane. (D) A representative Bioanalyzer (Agilent) lane profile showing the S1-seq library size distribution (predominantly 200–1000 bp).

###### 1.3.2 Embedding cells in plugs and deproteination

19. In a dry heating block, heat 1% low-melting-point agarose solution for 1-2 min at 95 °C. Place tubes at 50 °C after gently inverting and briefly spinning them to collect any liquid on the tube cap. Allow to equilibrate at 50 °C **≥**10 min. **<u>Note 3</u>:** Discard the tube if the cap opens during handling or heating because the final agarose concentration will be uncertain.
20. Remove the supernatant from tube in step 18 and resuspend cells with room temperature GBSS (without PVA or BSA). Add ∼73 μL GBSS first and then bring the final volume to 90 μl per two plugs. **<u>Note 4</u>:** Based on the cell and plug numbers that were determined in step 15, multiply the necessary volume accordingly.
21. To the room temperature cell suspension, add an equivalent amount of 50 °C 1% low-melting-point agarose. Mix by pipetting up and down. Fill each plug mold with ∼90 μl to slightly above the top, avoiding underfilling or overfilling (**Figure 6B**).
22. Solidify agarose plugs at 4 °C for 5 min and transfer two plugs into a 2 ml tube containing 1.5 ml of lysis buffer with 100 μg/ml of proteinase K. Tightly seal the tubes with paraffin film. Incubate at 50 °C while rotating slowly in a hybridization oven over two nights. Using a water bath or heating block instead is acceptable, but make sure the two plugs don’t stick to one another by gently inverting the tubes a few times during the incubation period.
23. Transfer the plugs into a 14 ml round-bottom tube after carefully pouring off the lysis buffer. It is okay to combine multiple plugs from the same sample in one 14 ml round bottom tube. **<u>Note 5</u>:** Plugs are now completely transparent and can be nearly invisible. Be careful not to discard the plugs when pouring the solution. Using a cell strainer could be helpful in preventing agarose plugs from falling into liquid waste.
24. Wash the plugs five times with 8 ml of TE, each time for 20 min at room temperature with gentle shaking. Check that no plugs are attached to the tube wall or cap.
25. After the last wash, discard as much TE as possible and add 1 ml of TE containing 100 μg/ml RNase A to the 14 ml round-bottom tube with plugs and incubate at 37 °C for 3 hr.
26. Wash plugs five times as in step 24.
27. After the last wash, replace with 8 ml of ice-cold TE and store plugs in the same 14 ml round-bottom tube at 4 °C until use. We found that it was normally acceptable to keep plugs for up to a month, but long-term storage may have an impact on library quality.

#### 2 Library preparation

##### 2.1 Key equipment and consumables

- Low-retention filter tips
- 2.0 ml low surface adhesion tubes
- 1.5ml DNA LoBind tubes (Effendorf)
- 14 ml round bottom tubes (Falcon, cat. no. 352059)
- 8-strip PCR tubes
- Stainless-steel lab spatula
- Microcentrifuge (conventional and refrigerated)
- Thermal mixer (e.g., Eppendorf ThermoMixer C)
- Thermal cycler
- Vortex mixer
- 0.22 μm cellulose nitrate sterilizing filter
- Water bath or cooling incubator set to 16 °C
- Covaris E220 Focused-ultrasonicator
- microTUBE-500 AFA Fiber Screw-Cap
- Qubit Fluorometer with dsDNA High Sensitivity (HS) assay kit (Invitrogen)
- Magnetic separation rack

##### 2.2 Reagents and buffers

- S1 nuclease (Promega, cat. no. M5761; 100 U/μl stock; Make 10 U/μl dilution with 1× S1 buffer freshly just before use)
- Exonuclease VII (NEB, cat. no. M0379L; 10 U/μl stock)
- Exonuclease T (NEB, cat. no. M0265L; 5 U/μl stock)
- NEBuffer 4 (NEB, cat. no. B7004S)
- Synthetic oligonucleotides (ligation adaptors and PCR primers; **Table 1**)
- β-agarase I (NEB, cat. no. M0392L)
- T4 DNA polymerase (NEB, cat. no. M0203L)
- T4 DNA ligase (NEB, cat. no. M0202T)
- SPRIselect beads (Beckman Coulter, cat. no. B23317; store at room temperature)
- AMPure XP beads (Beckman Coulter, cat. no. A63880; store at 4 °C)
- Dynabeads M-280 streptavidin (Thermo Fisher Scientific, cat. no.11206D; store at 4 °C)
- End-it DNA end-repair kit (Lucigen, cat. no. ER81050)
  - This product can be replaced by NEBNext End Repair Module (NEB, cat. no. E6050S).
- Phusion High-Fidelity DNA Polymerase (Thermo Fisher Scientific, cat. no. F503L)
- UltraPure DNase/RNase-free distilled water (Thermo Fisher Scientific, cat. no. 10977015)
- TE: 10 mM Tris-HCl pH 7.5, 1 mM EDTA
- 500 mM EDTA pH 8.0
- 10× S1 buffer: 500 mM sodium acetate, 2.8 M NaCl, 45 mM ZnSO_4_, pH 4.7 at 25 °C (filter sterilize and store at –20 °C; on each day of use, prepare 1× working solution with UltraPure water)
- 5× Exonuclease VII buffer: 250 mM Tris-HCl, 250 mM sodium phosphate, 40 mM EDTA, 50 mM 2-mercaptoethanol, pH 8 at 25 °C (filter sterilize and store at –20 °C; on each day of use, prepare 1× working solution with UltraPure water)
- 10× T4 ligase buffer (NEB, cat. no. B0202S)
- BSA (NEB, cat. no. B9000S) or recombinant albumin (NEB, cat. no. B9200S)
- dNTPs (Roche, cat. no. 11969064001)
- 1× T4 DNA polymerase buffer: 1× T4 ligase buffer containing 100 μg/ml BSA and 100 μM dNTPs (25 μM each)
- 10× β-agarase I buffer: 100 mM bis-Tris-HCl, 10 mM EDTA, pH 6.5 at 25 °C (filter sterilize and store at –20 °C; on each day of use, prepare 1× working solution with UltraPure water)
- Oligo dilution buffer: 10 mM Tris-HCl pH 7.5, 50 mM NaCl, 1 mM EDTA pH 8.0 (filter sterilize and store at room temperature; use to dissolve and dilute adaptors)
- 100% ethanol
- 3 M sodium acetate pH 5.5 (filter sterilize and store at –20 °C)
- 2× binding and washing (B&W) buffer: 10 mM Tris-HCl pH 7.5, 1 mM EDTA, 2 M NaCl (filter sterilize and store at room temperature; on each day of use, prepare 1× working solution with UltraPure water)
- 10 mM Tris-HCl, pH 7.5

**Table 1.**
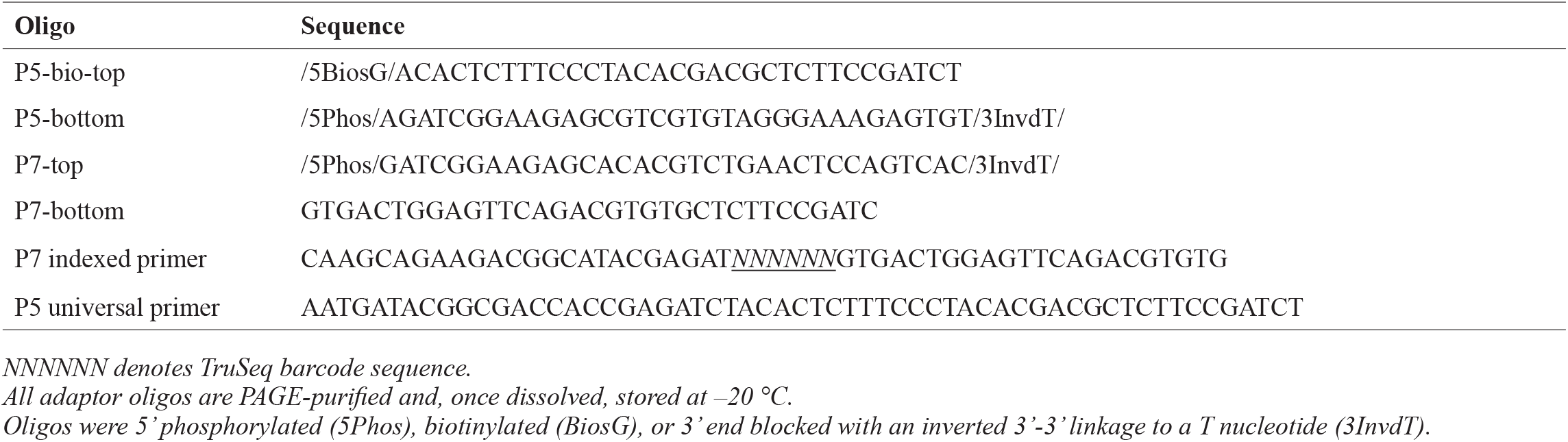
Oligonucleotide adaptors and sequencing primers.

##### 2.2 Procedures

###### Option 1. S1 nuclease treatment (for S1-seq)

1. Prepare 2 ml tubes with 500 μl TE. Place one plug in each tube. **<u>Note 6</u>**: If plugs were produced as described in section 1.3.2, one plug from juvenile mice (1.5∼2 million cells) and two plugs from adult mice (5∼6 million cells) should be used for each library preparation. If a 14 ml round-bottom tube contained more plugs than needed, drain the storage buffer (ice-cold TE) and use a stainless-steel spatula to carefully transfer the required number of plugs. To preserve the remaining plugs for future use, replenish the storage buffer.
2. Aspirate the TE and replace with 500 μl 1× S1 buffer. Equilibrate for 30 min at room temperature. **<u>Note 7</u>**: Use a 1 ml pipette and be careful not to damage the plug while aspirating or replacing buffers. It is critical to use home-made S1 buffer with pH 4.7–4.9 instead of pH 4.5 (as provided with the S1 enzyme).
3. Repeat step 2 three more times, for a total of 2 hr of equilibration.
4. Prepare 500 μl of S1 reaction solution (499.1μL × S1 buffer with 9 U of S1 nuclease; 100 U/μl stock, make a fresh 10 U/μl dilution with 1× S1 buffer just before use) per plug during the final equilibration, vortex to mix and keep on ice.
5. Aspirate liquid from the plugs and replace with 500 μl S1 reaction solution.
6. Place tubes on ice for 15 min to allow the enzyme to diffuse into the plugs. **<u>Note 8</u>:** This is important for homogeneous and uniform digestion of the DNA in the plugs.
7. Incubate at 37 °C for 20 min on a thermomixer at 400 rpm.
8. Inactivate S1 nuclease by addition of EDTA pH 8.0 to a final concentration of 10 mM and incubate on ice for 15 min.
9. Rinse plugs once with 500 μl ice-cold TE and continue to **Section 2.3.2**.

###### Option 2. Exonuclease VII treatment (for Exo7/T-seq)

1. Prepare 2 ml tubes with 500 μl of TE. Place one plug in each tube. **<u>Note 9</u>**: If plugs were produced as described in section 1.3.2, one plug from juvenile mice (1.5∼2 million cells) or two plugs from adult mice (5∼6 million cells) should be used for each library preparation. If a 14 ml round-bottom tube contains more plugs than needed, drain the storage buffer (ice-cold TE) and use a stainless-steel spatula to carefully transfer the required number of plugs. To preserve the remaining plugs for future use, replenish the storage buffer.
2. Aspirate TE buffer and replace with 1 ml 1× exonuclease VII buffer and equilibrate for 15 min at room temperature.
3. Repeat step 2 one more time, for a total of 30 min of equilibration.
4. Place samples on ice and incubate for 10 min. During the incubation, prepare 100 μl of exonuclease VII reaction solution (95 μl 1× exonuclease VII buffer with 50 U of exonuclease VII; 10 U/μl stock) per plug, vortex to mix and keep on ice.
5. Aspirate the liquid from the plugs and replace with 100 μl of exonuclease VII reaction solution. Incubate on ice for at least 20 min to allow the enzyme to diffuse into the plugs. **<u>Note 10</u>**: This is important for homogeneous and uniform digestion of the DNA in the plugs.
6. Incubate at 37 °C for 60 min on a thermomixer at 400 rpm.
7. Immediately place samples on ice after incubation.
8. Rinse plugs with 500 μl TE once and continue to **Option 3**, step2.

###### Option 3. Exonuclease T treatment (for Exo7/T-seq or Exo T-seq)

1. Prepare 2 ml tubes with 500 μl TE. Place one plug in each tube. **<u>Note 11</u>**: If plugs were produced as described in section 1.3.2, one plug from juvenile mice (1.5∼2 million cells) or two plugs from adult mice (5∼6 million cells) should be used for each library preparation. If a 14 ml round-bottom tube contained more plugs than needed, drain the storage buffer (ice-cold TE) and use a stainless-steel spatula to carefully transfer the required number of plugs. To preserve the remaining plugs for future use, replenish the storage buffer.
2. Aspirate the TE buffer from the plugs and replace with 1 ml 1× NEBuffer 4 and equilibrate for 15 min at 4 °C or on ice.
3. Repeat step 2 two more time, for a total of 45 min of equilibration.
4. Prepare 100 μl of exonuclease T reaction solution (85 μl of 1× NEB Buffer 4 containing 75 U of exonuclease T; 5 U/μl stock) per plug during the final equilibration, vortex to mix and keep on ice.
5. Aspirate the liquid from the plugs and replace with 100 μl of exonuclease T reaction solution and incubate on ice for 30 min to allow the enzyme to diffuse into the plugs. **<u>Note 12</u>**: This is important for homogeneous and uniform digestion of the DNA in the plugs.
6. Incubate at 24 °C for 90 min on a thermomixer at 400 rpm.
7. Immediately transfer the plugs into a 14 ml round-bottom tube after carefully aspirating off the exonuclease T reaction solution. It is okay to combine two plugs from the same sample in one 14 ml round-bottom tube if two plug were used from the same sample.
8. Wash the plugs five times with 8 ml of ice-cold TE, each time for 15 min at 4 °C with gentle shaking. Check that no plugs are attached to the wall or the cap.
9. After the last washing, transfer plugs to new 2 ml tubes with 500 μl ice-cold TE. The sample can be stored at 4 °C for a couple of days or before proceeding to **Section 2.3.2**.

###### T4 end filling

1. Equilibrate the plugs in 500 μl of 1× T4 DNA polymerase buffer for 30 min on ice.
2. Repeat step 1 three more times, for a total 2 hr of equilibration.
3. Prepare 500 μl of T4 DNA polymerase reaction solution (490 μl 1× T4 DNA polymerase buffer containing 30 U T4 DNA polymerase; 3 U/μl stock) per plug during the final equilibration, vortex to mix and keep on ice.
4. Aspirate the liquid from the plugs and replace with 500 μl of fresh T4 DNA polymerase reaction solution and incubate on ice for 15 min.
5. Incubation at 12 °C for 30 min on a thermomixer at 400 rpm. Occasionally tap the sample to ensure proper enzyme mixing.
6. The P5 adaptor (biotinylated and optionally nonbiotinylated for control) needs to be prepared fresh for use at step 12 below, so this is a good point to prepare it. Mix oligos P5-top (100 μM) and P5-bottom (100 μM) at equimolar concentration. Heat the mixture at 95 °C and allow to cool/anneal at room temperature for at least 1 hr (or use a thermocycler with the following program: 95 °C for 5 min; ramp down to 25 °C at 0.1 °C per second; 25 °C for 10 min; hold at 4 °C).
7. Inactivate the T4 DNA polymerase by addition of EDTA pH 8.0 to a final concentration of 10 mM, gently invert the tubes a couple of times, briefly spin down to ensure no plugs are stuck to the lids, and incubate on ice for 15 min.
8. Aspirate the liquid and replace with 1 ml TE, gently invert tubes, and incubate for 10 min on ice.
9. Repeat step 8 two more times.

###### Ligation of the first adaptor

10. Aspirate the TE from the plugs and replace with 500 μl of 1× T4 ligase buffer and incubate on ice for 15 min.
11. Repeat step 10 three more times, for a total of 1 hr of equilibration.
12. Aspirate the liquid and replace with 70 μl of 1× T4 ligase buffer containing 1 μl of the annealed P5 adaptor.
13. Equilibrate on ice for 15 min and add 1μl of T4 DNA ligase (2000 U/μl stock).
14. Incubate in a water bath at 16 °C for 20 hr or overnight.

###### Ligation buffer exchange

15. Spin down the reaction tube and aspirate and replace with 1 ml of ice-cold TE.
16. Immediately replace TE two more times.
17. Wash plugs three times with 1 ml of ice-cold TE on ice for 15 min each.
18. Transfer plugs to a 14 ml round-bottom tube (it is okay to combine two plugs in one 14 ml round bottom tube if you started with two plugs per sample) containing 8 ml of ice-cold TE and incubate at 4 °C for 15 min with gentle shaking.
19. Replace ice-cold TE and incubate overnight at 4 °C with gentle shaking.

###### Retrieval of DNA from plugs

1. Prepare 2 ml tubes with 500 μl TE and place one plug in each tube.
2. Aspirate TE and replace with 500 μl 1× β-agarase I buffer and equilibrate for 30 min at 4 °C or on ice.
3. Repeat step 2 one more time, for a total of 1 hr of equilibration.
4. Briefly spin down the tubes and aspirate as much buffer as possible.
5. Add 150 μl of 1× β-agarase I buffer.
6. Incubate at 70 °C for 5 min to melt the agarose.
7. Vortex for 30 sec and spin down.
8. Repeat steps 6 and 7 two more times.
9. Incubate at 42 °C for 5–10 min in a thermomixer.
10. During the incubation, prepare mixture of 8 μl of 1× β-agarase I buffer and 2 μl of β-agarase I per plug and keep at 42 °C.
11. Add 10 μl of β-agarase I solution from step 10 to each plug, mix well by tapping, and incubate at 42 °C for 1–1.5 hr with occasional vortexing and brief spinning (or vortex for 5 sec at 1400 rpm every 5 min on a thermomixer, spin down every 20 min to avoid condensation).
12. After the agarose is completely dissolved, use a P1000 pipette to measure the volume and add TE to bring to 500 μl. **<u>Note 13</u>:** The exact volume is critical because Covaris E220 sample chambers require 500 μl ± 5 μl for uniform shearing.

###### Shearing and precipitation of DNA

13. Transfer each dissolved sample to a microTUBE-500 with AFA Fiber and keep on ice.
14. Shear DNA into fragment sizes ranging 200–500 bp with a Covaris E220 Focused-ultrasonicator using the following parameters: temperature 7 °C; instrument setting as delay 300 sec then three iterations of [peak incident power 175 W, duty factor 20%, cycles/burst 200, duration 30 sec, and delay 90 sec].
15. Prepare new 2 ml tubes with 1250 μl of 100% ethanol for each sample.
16. Save 5 μl of sonicate samples to check shearing by agarose gel electrophoresis (**Figure 6C**).
17. Transfer the remainder of each sonicated sample to a 2 ml tube containing ethanol, add 55 μl of 3 M sodium acetate, and incubate overnight at –20 °C.
18. Spin the samples at maximum speed at 4 °C for 15 min in a microcentrifuge. Turn the tubes around 180° and spin for another 5 min so that the DNA pellet is collected properly on one side of the 2 ml tube. **<u>Note 14</u>**: If this step is not followed as described, the DNA will be smeared along the tube’s length on one side.
19. Wash the pellet with 500 μl ice-cold 70% ethanol (freshly prepared) and centrifuge for 5 min at 4 °C.
20. Aspirate the ethanol and air dry the pellet for 15 min at room temperature.
21. Add 50 μl of TE and incubate 1 hr at 37 °C to dissolve the DNA pellet.
22. Measure the DNA concentration with Qubit, dsDNA HS assay kit.

###### Size selection (removal of unligated adaptor)

23. Before use, mix SPRIselect beads well by vortexing for 30 sec.
24. Transfer the dissolved DNA samples to new 1.5 ml DNA LoBind tubes. If you started with two plugs per sample, combine the dissolved DNA samples at this point.
25. Add 1.2× sample volumes of SPRIselect beads (60 μl if you started with one plug; 120 μl if two plugs) to each sample to select DNA above 300 bp, which will exclude the free adaptor DNA.
26. Mix well by pipetting up and down ten times and incubate at room temperature for 5 min.
27. Place the tubes on a magnetic separation rack and wait until the solution is completely clear.
28. Remove the supernatant. **<u>Note 15</u>:** Do not try to remove all the liquid as this will remove some beads as well.
29. Keep the tubes on the magnetic rack and add 200 μl of freshly prepared 70% ethanol and turn the tubes to the other side to allow the beads to wash. Turn the tubes back and aspirate all of the wash buffer.
30. Repeat the 70% ethanol wash once and remove the wash buffer completely.
31. Let the beads air dry for 10 min or until tiny cracks start to appear. **<u>Note 16</u>:** It is important to dry the beads completely to avoid carry-over of ethanol to the next steps.
32. Add 42 μl of TE to elute the DNA, remove the tubes from the magnetic rack, thoroughly resuspend the beads by pipetting with a P200 pipette and incubate for at least 10 min at room temperature.
33. Place the tubes on the magnetic rack and transfer 40 μl of the eluate to a new 1.5 ml DNA LoBind tube, ensuring that there are no SPRIselect beads carried over.

###### Streptavidin pulldown of biotinylated adaptor-ligated DNA

1. Prepare 2× and 1× B&W buffers.
2. Calculate the amount of Dynabeads M-280 Streptavidin required: 20 μl per library sample.
3. Vortex the Dynabeads thoroughly before use, then transfer the entire volume needed into a single 1.5 ml DNA LoBind tube.
4. Wash the beads by adding 1 ml of 1× B&W buffer and vortexing for 5 sec.
5. Place the tube on a magnetic rack for 1 min or until the solution is completely cleared. Discard the supernatant.
6. Remove the tube from the magnetic rack and resuspend the beads in 100 μl of 1× B&W buffer per sample, plus 50 μl extra. (i.e., if you have four samples, resuspend in 450 μl.) Mix well by pipetting.
7. For each sample, aliquot 100 μl of the bead suspension into a fresh 1.5 ml DNA LoBind tube.
8. To the remaining bead suspension in the wash tube, add a further 100 μl of 1× B&W buffer per sample and mix well by pipetting. Transfer 100 μl of this bead suspension into each of the tubes in step 7. The total volume should be 200 μl for each tube.
9. Place the tubes on the magnetic rack and carefully aspirate the supernatant completely.
10. Remove the tubes from the magnetic rack and resuspend the beads with 40 μl of 2× B&W buffer.
11. Place the samples on the magnetic separation rack for 3 min to remove any remaining SPRIselect beads.
12. Carefully transfer 40 μl of the DNA sample to the washed Dynabeads from step 10. Mix well by pipetting.
13. Incubate at room temperature for 30 min on a roller. Gently vortex the samples every 10 min to prevent the beads from settling down. (Optional: mix for 10 sec at 1400 rpm every 10 min on a thermomixer at 24 °C).
14. Place the tubes on the magnetic rack. Turn the tubes around a few times in the rack to collect all of the beads on one side of the tube. If necessary, use a P1000 pipette to pipette slowly once or twice to collect beads stuck on the wall.
15. Wash the beads with 500 μl of 1× B&W buffer three times. For each wash, mix the beads well by vortexing for 5 sec, use a quick spin to collect the liquid, and return the tube to the magnetic rack. Wait until the solution is completely cleared and aspirate the supernatant.
16. Wash the beads with 500 μl of 10 mM Tris-HCl pH 7.5 two times in the same way as in step 15.
17. During the last wash, prepare the 1× end-repair reaction mix, using the End-it DNA End-repair kit (**Table 2.1**.) or 1× end repair module mix, using the NEBNext End Repair Module (**Table 2.2**.). The reaction volume needed per sample is 100 μl. Scale accordingly.
18. Resuspend the beads in 100 μl 1× end-repair mix.
19. If using the End-it DNA End-repair kit, incubate at room temperature for 45 min on a roller. Gently vortex the samples every 10 min to prevent the beads from settling down. (Optional: mix for 10 sec at 1400 rpm every 10 min on a thermomixer at 24 °C.) If using the NEBNext End Repair Module, incubate at 20 °C for 30 min on a roller or on a thermomixer.
20. At this point, prepare the P7 adaptor by mixing oligos P7-top and P7-bottom at equimolar concentration (100 μM each). Heat the mixture at 95 °C and allow to cool/ anneal at room temperature for at least 1 hr. (Optional: use a thermocycler with following profile: 95 °C for 5 min; ramp down to 25 °C at 0.1 °C per sec; 25 °C for 10 min; hold at 4 °C).
21. Capture the beads on the magnetic rack and wash with 500 μl of 1× B&W buffer three times in the same way as in step 15.
22. Wash the beads two times with 500 μl of 10 mM Tris-HCl pH7.5.
23. Wash the beads with 500 μl of 1× T4 DNA ligase buffer.
24. Wash the beads with 100 μl of 1× T4 DNA ligase buffer. Ensure that all of the beads sticking to the walls of the tube are completely resuspended.
25. Prepare the ligation reaction mixture (**Table 3**).
26. Use the magnetic rack to capture the beads and remove the supernatant. Resuspend the beads in 98 μl of the ligase reaction mixture and place on ice.
27. Add 2 μl of the annealed P7 adaptor.
28. Incubate at 16 °C on a roller for 20 hr or overnight.

**Table 2.1.** 1 × end-repair reaction mix.

|  |  |
| --- | --- |
| UltraPure water | 68 µl |
| 10× Buffer | 10 µl |
| dNTPs | 10 µl |
| ATP | 10 µl |
| Enzyme Mix | 2 µl |

**Table 2.2.** 1 × end repair module mix.

|  |  |
| --- | --- |
| UltraPure water | 85 µl |
| 10× Buffer | 10 µl |
| Enzyme Mix | 5 µl |

**Table 3.** 1 × ligation reaction mix.

|  |  |
| --- | --- |
| UltraPure water | 87.8 µl |
| 10× T4 DNA Ligase Buffer | 10 µl |
| T4 DNA Ligase (2000 U/µl) | 0.2 µl |

###### PCR amplification

1. Capture the beads on a magnetic rack and wash two times with 500 μl of 1× B&W buffer.
2. Resuspend the beads in 500 μl of 1× B&W buffer and transfer to a new 1.5 ml DNA LoBind tube.
3. Wash the beads two times with 500 μl of 10 mM Tris-HCl pH 7.5.
4. Resuspend the beads in 20 μl of 10 mM Tris-HCl pH 7.5. Make sure that the resuspension is uniform and that there are no beads stuck to the tube walls.
5. Divide each 20 μl of bead suspension into two PCR tubes and prepare 1× PCR reaction mix (**Table 4**).
6. Run the PCR with the following conditions: 98 °C for 30 sec; 16 cycles of 98 °C for 10 sec, 65 °C for 20 sec, and 72 °C for 20 sec; and a final extension step at 72 °C for 5 min.

**Table 4.** 1 × PCR reaction mix.

|  |  |
| --- | --- |
| Beads in 10 mM Tris-HCl pH 7.5 | 10 µl |
| UltraPure water | 24.5 µl |
| 5× Phusion HF Buffer | 10 µl |
| 10 µM Primer P5 | 2 µl |
| 10 µM Primer P7 (indexed) | 2 µl |
| 10 mM dNTPs | 1 µl |
| Phusion Polymerase (2 U/µl) | 0.5 µl |

###### Size selection using AMPure XP beads

1. Handle PCR products on a different bench from the one used to set up the reaction and use designated pipettes and tips, as well as reagent aliquots reserved for PCR products. UV-irradiate both side of the pipettes and the magnetic rack using 1200 J/m^2^ before use. These steps are critical to avoid cross-contamination of sequencing libraries.
2. During the PCR, remove the AMPure XP beads from 4 °C storage and keep at room temperature for 30 min. Mix well by vortexing for 30 sec just before use.
3. For each library sample, combine both PCRs in a new 1.5 ml DNA LoBind tube.
4. Add 0.9× sample volumes of AMPure bead suspension to each sample and mix by pipetting up and down at least 10 times.
5. Incubate at room temperature for 8 min.
6. Place the tubes on a magnetic rack for 20 min or until the solution is completely clear.
7. Remove the supernatant. Do not try to remove all the solution as this will remove some beads as well.
8. Keep the tubes on the magnetic rack and add 200 μl of freshly prepared 80% ethanol without disturbing the beads, to wash away contaminants. Wait for 30 sec, then carefully remove the supernatant.
9. Repeat step 8 one more time.
10. Let the beads air dry for 12 min.
11. Remove tubes from the magnetic rack and resuspend the beads in 20 μl of 10 mM Tris-HCl pH 7.5. Mix thoroughly by pipetting and incubate for at least 7 min at room temperature.
12. Place the tubes on the magnetic rack for 5 min or longer until the solution is completely clear.
13. Collect the clear supernatant to a new 1.5 ml DNA Lo-Bind tube.
14. To verify the size range of the library, a dilution of the library is analyzed on a Bioanalyzer Instrument (Agilent Technologies) according to the manufacturer’s instructions. An example is shown in **Figure 6D**.
15. Proceed with next-generation sequencing (50 bp pair-end reads) or store at –20 °C until ready to sequence.

#### 3 Bioinformatic analysis

Sequence reads are trimmed and filtered by Trim Galore (http://www.bioinformatics.babraham.ac.uk/projects/trim_galore/) with the arguments --paired --length 15. Reads are mapped onto the mouse reference genome (mm10) using bowtie2 (Langmead et al. 2009) with the arguments -N 1 -X 1000. Duplicated reads are removed using Picard (https://broadinstitute.github.io/picard/). Uniquely and properly mapped reads (MAPQ **≥** 20) are extracted using samtools (Danecek et al. 2021) with the argument -q 20. Reads are counted at the nucleotide next to where the biotinylated adaptor DNA was mapped (this corresponds to the position of the respective resection endpoint). Code used for read processing and mapping is available online at https://github.com/yamadas2/mouse-S1seq, which also includes scripts for generating mean profiles and heatmaps around SPO11-oligo hotspot centers, and scatter plots to check correlation with SPO11-oligo maps and reproducibility between maps. New sequencing data have been deposited in the Gene Expression Omnibus (GEO) repository under accession number GSE265863.

## Acknowledgments

This article is subject to the Open Access to Publications policy of the Howard Hughes Medical Institute (HHMI). HHMI lab heads have previously granted a nonexclusive CC BY 4.0 license to the public and a sublicensable license to HHMI in their research articles. Pursuant to those licenses, the author-accepted manuscript of this article can be made freely available under a CC BY 4.0 license immediately upon publication. We thank Eleni Mimitou and Pei-Ching Huang for their contributions to the development (E.M.) and application (E.M. and P.C.H.) of S1-seq to yeast meiosis. We thank Neeman Mohibullah and the MSK Integrated Genomics Operation (IGO) for sequencing and the MSK Mouse Genetics Colony Management Group for assistance with mouse husbandry. MSK core facilities are supported by the US National Cancer Institute cancer center support grant P30 CA08748. The IGO was further funded by the Cycle for Survival and the Marie-Josée and Henry R. Kravis Center for Molecular Oncology. This work was supported by US National Institutes of Health grant R35 GM118092 (to S. Keeney) and the Brain Pool Program through the National Research Foundation of Korea (NRF) funded by the Ministry of Science and ICT (2022H1D3A2A01096332 to S. Kim).

## References

Amente S, Scala G, Majello B, Azmoun S, Tempest HG, Premi S, Cooke MS. 2021. Genome-wide mapping of genomic DNA damage: methods and implications. Cell Mol Life Sci 78: 6745–6762.

Bellve AR, Cavicchia JC, Millette CF, O’Brien DA, Bhatnagar YM, Dym M. 1977. Spermatogenic cells of the prepuberal mouse. Isolation and morphological characterization. J Cell Biol 74: 68–85.

Bergerat A, de Massy B, Gadelle D, Varoutas PC, Nicolas A, Forterre P. 1997. An atypical topoisomerase II from Archaea with implications for meiotic recombination. Nature 386: 414–417.

Canela A, Maman Y, Jung S, Wong N, Callen E, Day A, Kieffer-Kwon KR, Pekowska A, Zhang H, Rao SSP et al. 2017. Genome organization drives chromosome fragility. Cell 170: 507–521 e518.

Canela A, Sridharan S, Sciascia N, Tubbs A, Meltzer P, Sleckman BP, Nussenzweig A. 2016. DNA Breaks and End Resection Measured Genome-wide by End Sequencing. Mol Cell 63: 898–911.

Cejka P, Symington LS. 2021. DNA End Resection: Mechanism and Control. Annu Rev Genet 55: 285–307.

Chase JW, Richardson CC. 1974a. Exonuclease VII of Escherichia coli. Mechanism of action. J Biol Chem 249: 4553–4561.

Chase JW, Richardson CC. 1974b. Exonuclease VII of Escherichia coli. Purification and properties. J Biol Chem 249: 4545–4552.

Claeys Bouuaert C, Tischfield SE, Pu S, Mimitou EP, Arias-Palomo E, Berger JM, Keeney S. 2021. Structural and functional characterization of the Spo11 core complex. Nat Struct Mol Biol 28: 92–102.

Danecek P, Bonfield JK, Liddle J, Marshall J, Ohan V, Pollard MO, Whitwham A, Keane T, McCarthy SA, Davies RM et al. 2021. Twelve years of SAMtools and BCFtools. Gigascience 10.

Deutscher MP, Marlor CW. 1985. Purification and characterization of Escherichia coli RNase T. J Biol Chem 260: 7067–7071.

Gittens WH, Johnson DJ, Allison RM, Cooper TJ, Thomas H, Neale MJ. 2019. A nucleotide resolution map of Top2-linked DNA breaks in the yeast and human genome. Nature communications 10: 4846.

Guillon H, Baudat F, Grey C, Liskay RM, de Massy B. 2005. Crossover and noncrossover pathways in mouse meiosis. Molecular Cell 20: 563–573.

Huang SN, Michaels SA, Mitchell BB, Majdalani N, Vanden Broeck A, Canela A, Tse-Dinh YC, Lamour V, Pommier Y. 2021. Exonuclease VII repairs quinolone-induced damage by resolving DNA gyrase cleavage complexes. Sci Adv 7.

Hunter N. 2015. Meiotic Recombination: The Essence of Heredity. Cold Spring Harb Perspect Biol 7.

Keeney S, Giroux CN, Kleckner N. 1997. Meiosis-specific DNA double-strand breaks are catalyzed by Spo11, a member of a widely conserved protein family. Cell 88: 375–384.

Lange J, Pan J, Cole F, Thelen MP, Jasin M, Keeney S. 2011. ATM controls meiotic double-strand-break formation. Nature 479: 237–240.

Langmead B, Trapnell C, Pop M, Salzberg SL. 2009. Ultrafast and memory-efficient alignment of short DNA sequences to the human genome. Genome Biol 10: R25.

Liu C, Hauk G, Yan Q, Berger JM. 2024. Structure of Escherichia coli exonuclease VII. Proc Natl Acad Sci U S A 121: e2319644121.

Liu J, Wu TC, Lichten M. 1995. The location and structure of double-strand DNA breaks induced during yeast meiosis: evidence for a covalently linked DNA-protein intermediate. EMBO J 14: 4599–4608.

Maekawa K, Yamada S, Sharma R, Chaudhuri J, Keeney S. 2022. Triple-helix potential of the mouse genome. Proc Natl Acad Sci U S A 119: e2203967119.

Mimitou EP, Keeney S. 2018. S1-seq Assay for Mapping Processed DNA Ends. Methods Enzymol 601: 309–330.

Mimitou EP, Yamada S, Keeney S. 2017. A global view of meiotic double-strand break end resection. Science 355: 40–45.

Mirkin SM, Frank-Kamenetskii MD. 1994. H-DNA and related structures. Annu Rev Biophys Biomol Struct 23: 541–576.

Paiano J, Wu W, Yamada S, Sciascia N, Callen E, Paola Cotrim A, Deshpande RA, Maman Y, Day A, Paull TT et al. 2020. ATM and PRDM9 regulate SPO11-bound recombination intermediates during meiosis. Nature communications 11: 857.

Pan J, Sasaki M, Kniewel R, Murakami H, Blitzblau HG, Tischfield SE, Zhu X, Neale MJ, Jasin M, Socci ND et al. 2011. A hierarchical combination of factors shapes the genome-wide topography of yeast meiotic recombination initiation. Cell 144: 719–731.

Symington LS. 2016. Mechanism and regulation of DNA end resection in eukaryotes. Crit Rev Biochem Mol Biol 51: 195–212.

Wong N, John S, Nussenzweig A, Canela A. 2021. END-seq: An Unbiased, High-Resolution, and Genome-Wide Approach to Map DNA Double-Strand Breaks and Resection in Human Cells. Methods Mol Biol 2153: 9–31.

Yamada S, Hinch AG, Kamido H, Zhang Y, Edelmann W, Keeney S. 2020. Molecular structures and mechanisms of DNA break processing in mouse meiosis. Genes Dev 34: 806–818.

Zelazowski MJ, Sandoval M, Paniker L, Hamilton HM, Han J, Gribbell MA, Kang R, Cole F. 2017. Age-dependent alterations in meiotic recombination cause chromosome segregation errors in spermatocytes. Cell 171: 601–614 e613.

Zuo Y, Deutscher MP. 1999. The DNase activity of RNase T and its application to DNA cloning. Nucleic Acids Res 27: 4077–4082.

